# The polyamines spermine and spermidine inhibit or induce programmed cell death in *Arabidopsis thaliana in vitro* and *in vivo* in a dose dependent manner

**DOI:** 10.1101/2023.11.15.567161

**Authors:** Rory Burke, Daniele Nicotra, Jim Phelan, Paul F. McCabe, Joanna Kacprzyk

**Affiliations:** School of Biology and Environmental Science, University College Dublin, Dublin 4, Ireland; Department of Agriculture, Food and Environment, University of Catania, 95123 Catania, Italy

## Abstract

Polyamines are ubiquitous biomolecules with a number of established functions in eukaryotic cells. In plant cells, polyamines have previously been linked to abiotic and biotic stress tolerance, as well as to the modulation of programmed cell death (PCD), with contrasting reports on their pro-PCD and pro-survival effects. Here, we used two well established platforms for the study of plant PCD; *Arabidopsis thaliana* suspension cultures cells and the root hair assay, to examine the roles of the polyamines spermine and spermidine in the regulation of PCD. We demonstrate that both polyamines can trigger PCD when applied exogenously at higher doses, whereas at lower concentrations they inhibit PCD induced by both biotic and abiotic stimuli. Furthermore, we show that concentrations of polyamines resulting in inhibition of PCD generated a transient ROS burst in our experimental system, and activated the expression of oxidative stress- and pathogen response-associated genes. Finally, we examined PCD responses in existing *Arabidopsis* polyamine synthesis mutants, and identified a subtle PCD phenotype in *Arabidopsis* seedlings deficient in thermo-spermine. The presented data show that polyamines can have a role in PCD regulation, however that role is dose-dependent and consequently they may act as either inhibitors, or inducers, of PCD in *Arabidopsis*.

## Introduction

Polyamines (PAs) are small organic nitrogenous bases present across all domains of life (Salvi and Tavladoraki, 2020). Since their identification (Bachrach, 2010), PAs have been linked to the regulation of a large number of biological processes in eukaryotes (Chen et al., 2019; Pegg, 2016). In plants, the most abundant PAs are spermine (SPM), spermidine (SPD) and putrescine (PUT), with thermospermine and cadaverine also present in some species and tissues (Minocha et al., 2014; Jancewicz et al., 2016). The biosynthetic pathways of plant PAs are well established (reviewed in Chen et al., 2019). Briefly, PA biosynthesis begins with the conversion of arginine to PUT, which can then be converted to SPD by spermidine synthase (SPDS) and then to SPM by spermine synthase (SPMS) or to thermospermine by ACL5, while back conversion of SPM to SPD to PUT is mediated by polyamine oxidase (PAO), generating H_2_O_2_. In animals, PAs appear to play an important role as regulators of programmed cell death (PCD) (Thomas* and Thomas, 2001), a genetically encoded mechanism of organised self-destruction of damaged or redundant cells, that needs to be tightly controlled for the benefit of the whole organism (Ellis Hm Fau - Horvitz and Horvitz, 1986; Danon et al., 2000; Ameisen, 2002). PA-mediated regulation of mammalian apoptosis appears complex, with both PAs accumulation and depletion having been linked to both the induction and inhibition of cell death pathways (Seiler and Raul, 2005). Many causative mechanisms have been suggested, including, but not limited to, interaction with apoptotic signalling cascades (Bratton et al., 1999; Bhattacharya et al., 2003), inhibition of endonuclease enzymes (Brüne et al., 1991) and both scavenging and production of cytotoxic reactive oxygen species (ROS) (Yoda et al., 2003; Chaturvedi et al., 2004). The well-established role of PAs in both cell proliferation and cell death in mammals is highlighted by emerging anti-cancer strategies based on targeting PA metabolism (Casero et al., 2018).

In plants, carefully controlled PCD events are an essential part of development, abiotic stress responses and host-pathogen interactions (Greenberg and Yao, 2004; Petrov et al., 2015; Daneva et al., 2016; Burke et al., 2020). While our understanding of plant PCD signalling is still fragmented in comparison to the well-characterized animal cell death pathways, there is substantial evidence suggesting that PAs may contribute to finely tuned regulation of cell death in plants via mechanisms both analogous to, and distinct from, those identified in animals (reviewed by Moschou and Roubelakis-Angelakis, 2014). For example, PAs appear to be involved in regulation of the hypersensitive response (HR), a rapid form of localized cell death induced to restrict the growth of invading pathogens (Morel and Dangl, 1997). PAs accumulate during HR induced by tobacco mosaic virus in *Nicotiana tabacum* (Torrigiani et al., 1997), and PA degradation, yielding hydrogen peroxide, appears to be necessary for the oxidative burst, an essential component of HR associated PCD (Yoda et al., 2003; Yoda et al., 2006). Additionally, SPM has been shown to induce the expression of several HR marker genes downstream of ROS signalling, Ca^2+^ influx and mitochondrial disruption, all of which have previously been associated with various forms of plant PCD (Takahashi et al., 2003). Indeed, transcriptomic analyses indicated that in *Arabidopsis*, exogenous SPM triggers expression of a set of genes, similar to those induced during HR, in response to cucumber mosaic virus (CMV), consequently increasing resistance to CMV infection (Mitsuya et al., 2009). PUT can trigger salicylic acid (SA) accumulation, and expression of SA associated defence genes required for ROS-dependent systemic acquired resistance (SAR) in *Arabidopsis* (Liu et al., 2020). PAs can also influence plant responses to abiotic stress (Gill and Tuteja, 2010). While most studies focus on general plant stress tolerance, and do not examine the influence of PAs on the fate of individual cells, there are studies showing that they may modulate plant cell death or survival in response to abiotic stress. For example, tobacco plants overexpressing PAO are more sensitive to PCD induced by methyl viologen and menadione (Moschou et al., 2008a). Likewise, plants with reduced apoplastic PAO activity demonstrated reduced cell death induced by salt stress (Moschou et al., 2008b). A similar effect was observed in cucumber where exogenous SPD application caused H_2_O_2_ accumulation, triggered transcription of ROS responsive genes and improved resistance to salt stress (Wu et al., 2018). Moreover, inhibition of PAO activity, or addition of PUT, reduced cell death in wheat root tips exposed to aluminium stress (Yu et al., 2018), and cadmium stress induced accumulation of PAs before PCD induction, in both *Malus hupehensis* seedlings (Jiang et al., 2012) and in tobacco BY-2 cells (Kuthanová et al., 2004), suggesting a role for PAs in regulation of cell death induced by heavy metal stress. PAs have also been associated with several forms of developmental PCD in plants. In *Arabidopsis*, *acl5* mutants defective in thermospermine synthesis (Knott et al., 2007) display perturbed xylem vessel development due to premature cell death in the developing vessels (Muñiz et al., 2008). The *acl5* xylem vessel phenotype could be reduced via supplementation of exogenous SPM, or replicated via induction of another form of cell death in the vessel elements, suggesting that the role of ACL5 in this context is to prevent premature cell death in the developing xylem vessels (Muñiz et al., 2008). Moreover, exogenous PAs supplementation has long been known to delay senescence (Cohen et al., 1979). In dark-induced barley leaf senescence, a process characterised by gradual organellar breakdown culminating in PCD, PAs accumulate during the early stages before decreasing immediately prior to cell death as the transcription of PA catabolism genes peaks (Sobieszczuk-Nowicka et al., 2016). Interestingly, while chemical inhibition of PUT oxidation accelerated senescence, inhibition of SPM and SPD oxidation had the opposite effect on the process, suggesting that different PAs may act to regulate senescence and the associated cell death pathway via separate mechanisms (Sobieszczuk-Nowicka et al., 2016).

In conclusion, while there exists extensive evidence linking PAs to modulation of plant PCD in numerous cell death contexts and species, the details of this connection require further elucidation. For example, the reported effects of exogenous PAs are often conflicting, and it appears that PAs are capable of pro-survival or pro-death roles in different developmental or environmental contexts. Likewise, many studies exploring the effect of PA treatment on plant tissues often suffer from a lack of specificity to conclusively record a bona fide genetically controlled cell death programme, rather than sub-lethal stress in cells that don’t die or treatments causing uncontrolled, accidental cell death known as necrosis (Reape et al., 2008; Kacprzyk et al., 2017). Additionally, studies of the effects of PAs on plant stress tolerance in many cases do not offer temporal or spatial resolution allowing quantification of cells undergoing PCD. Here, we used two well-established models for quantitative determination of plant PCD rates; *Arabidopsis* cell suspension cultures (ACSC) (McCabe and Leaver, 2000; Reape et al., 2008; Burke et al., 2023) and the root hair assay (RHA) (Hogg et al., 2011), to investigate the link between chemical and genetic perturbation of PA levels and PCD induced by biotic-like (salicylic acid) and abiotic (salinity) stimuli in a dose dependent manner. We also demonstrate that exogenous PAs induce ROS in *Arabidopsis* roots and activate transcription of oxidative stress responsive genes in cell culture, supporting the hypothesis that H_2_O_2_ from PA catabolism plays a key role in the mechanism of PA mediated regulation of PCD in plants. We also uncover a subtle, stress induced PCD phenotype for the thermo-spermine synthesis mutant *alc5,* that was previously used to demonstrate the role of PAs in abiotic stress responses (Yamaguchi et al., 2006; Kusano et al., 2007; Alet et al., 2012; Zarza et al., 2017). Collectively, the presented data further informs our understanding of the complex relationship between PAs and cell fate decisions in plants.

## Results

### Spermine and spermidine induce PCD in Arabidopsis thaliana ACSCs and root hair cells

Accumulation of PAs was demonstrated to induce apoptosis in some mammalian cell lines (Poulin et al., 1995; Seiler and Raul, 2005); however, the ability of exogenous PAs to induce PCD in individual plant cells has not yet been demonstrated. Similarly, while exogenous PAs have been shown to cause tissue damage in certain plant tissues (Yoda et al., 2003) the nature of this damage and whether or not it occurs via a cell death programme or uncontrolled necrosis was not investigated. Here, the ACSCs, that offers an established system for studying plant PCD, was treated with the PAs SPM and SPD (1 µM – 10 mM) and resulting rates of viability, PCD and necrosis determined after 24 hours (Figure 1). High concentrations (10 mM) of both PAs caused significant viability loss, with the majority of dying cells undergoing PCD, with characteristic protoplast retraction (Figure 1A), and only a small number of cells undergoing necrosis (Figure 1B). Five-day old *Arabidopsis* seedlings were used to demonstrate a similar effect of exogenous PAs *in planta*. Seedlings were treated with SPM and SPD (10 µM – 1 mM) and the root hairs scored as viable, PCD or necrotic using light microscopy after 24 hours using the root hair assay (Hogg et al., 2011; Kacprzyk et al., 2014). Both SPM and SPD induced a significant reduction in viability, with the majority of root hairs dying via PCD (Figure 1C).

**Figure 1.**
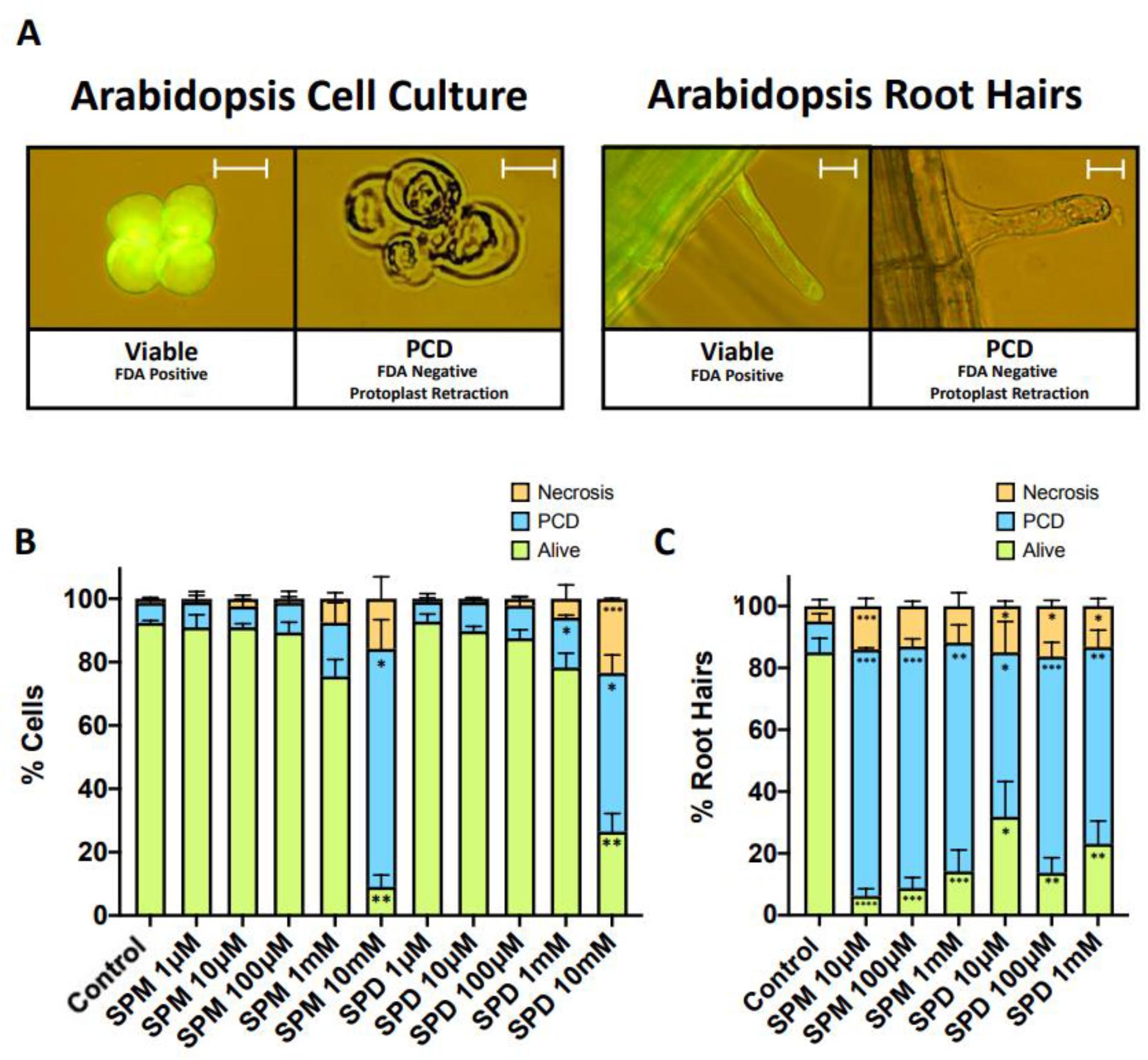
High concentrations of polyamines induce PCD in both ACSCs and *Arabidopsis* root hairs. Characteristic PCD morphology induced in ASCS and root hairs by PA treatment **(A).** ACSCs **(B)** and seedlings **(C)** were treated with varying concentrations of SPM and SPD and scored after 24 hours for viability and PCD or necrotic morphology. Experiments were repeated 3 **(B)** and 4 **(C)** times. Rates of alive, PCD and necrotic cells in each PA treatment were compared to control paired using 2-way ANOVA: *≤0.05, **≤0.01, ***≤0.001, ****≤0.0001. Error bars = SEM.

### Pre-treatment of ACSCs and Arabidopsis seedlings with exogenous polyamines protects against PCD induced by the plant hormone salicylic acid

There is extensive evidence linking PAs to PCD regulation (Moschou and Roubelakis-Angelakis, 2014), and exogenous PA supplementation to increased stress tolerance in plants (Gill and Tuteja, 2010). To examine if PAs could inhibit PCD in this experimental system, the ACSCs were pre-treated with SPM and SPD (500 µM, 1 mM, 5 mM) for 24 hours before PCD induction with salicylic acid (SA). SA is a plant hormone involved in pathogen induced PCD, such as the hypersensitive response (HR) (Heath, 2000; Radojičić et al., 2018), and can thus be used to mimic PCD occurring in a biotic context (Burke et al., 2023). Additionally, PAs have also previously been implicated in the modulation of HR progression (Yoda et al., 2003). The concentration of SA used (1.5mM) induced a high level of PCD in ACSCs, without significant impact on the rates of uncontrolled necrosis (Figure 2A). While the highest concentration of SPM (5 mM) induced PCD in control samples (Figure 2A), it also blocked PCD induced by SA treatment, as did pre-treatment with 5 mM SPD (Figure 2A). Moreover, pre-treatment with 1 mM SPM and SPD and 500 µM SPM reduced PCD rates of SA treated cells without significant toxic effect in control samples: in all cases, levels of necrosis remained low and induction or inhibition of PCD was thus associated with a reduction or increase in cell viability respectively (Figure S1A). The results obtained using the RHA recapitulated these observations *in planta.* Five-day old *Arabidopsis* seedlings were pre-treated in 100 nM or 1 µM SPM/SPD for 24 hours before PCD induction with 20 µM SA, and resulting rates of viability, PCD and necrosis were determined after 24h. Both tested concentrations of SPM and SPD had only minor effects on PCD rates of root hairs in control seedlings (absence of SA treatment), while 100 nM SPM, 100 nM SPD, 1 µM SPM and 1 µM SPD significantly inhibited PCD induction in root hair cells in seedlings treated with SA (Figure 2B). As with the ACSCs, inhibition of PCD in root hairs by SPM and SPD resulted in significant increases in viability, rather than necrosis (Figure S1B).

**Figure 2.**
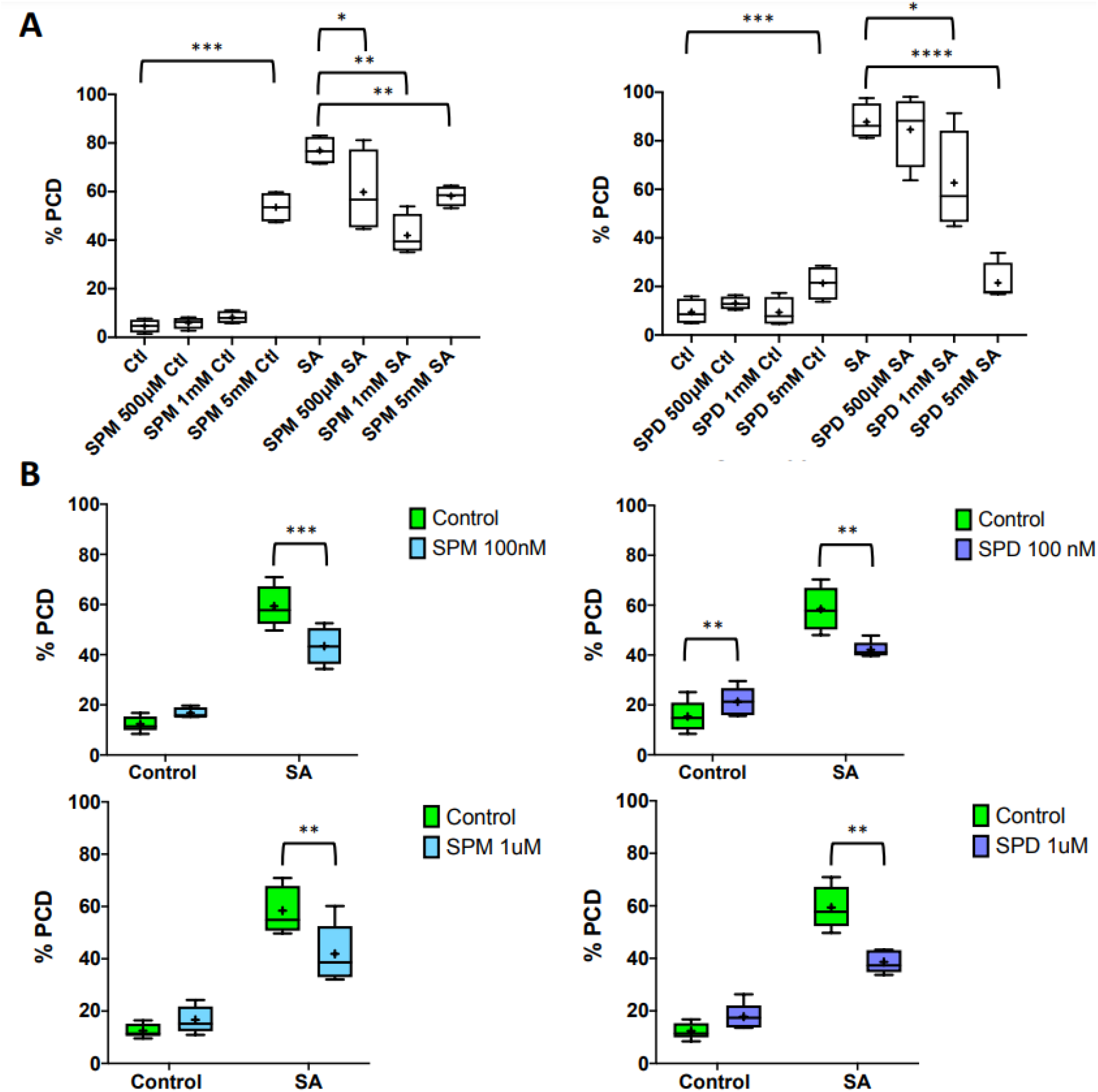
Pre-treatment with SPM or SPD protects ACSCs and *Arabidopsis* root hairs against SA-induced PCD. ACSCs were treated with indicated concentrations of SPM or SPD for 24 h prior to PCD induction with 1.5mM SA **(A)**. Cells were scored as viable, PCD or necrotic 24 h later. Experiments were repeated 5 times and rates of PCD cells in each treatment compared to control using 1-way ANOVA. *Arabidopsis* seedlings were treated with 100 nM SPM, 100 nM SPD, 1 µM SPM or 1 µM SPD for 24 hours, before PCD induction with 20 µM SA **(B)**. Root hairs were then scored as viable, PCD or necrotic 24 hours later. For each treatment 3 seedlings were examined per experiment, and experiments were repeated 5 times. Rates of PCD root hairs in each treatment were compared to control using 1-way ANOVA. *≤0.05, **≤0.01, ***≤0.001, ****≤0.0001. Lines on box and whisker plots show median values, crosses show mean values and whiskers show the minimum to maximum values.

### Arabidopsis seedlings pre-treated with exogenous SPM and SPD show increased tolerance against NaCl-induced PCD

To establish if the protective effect of PAs was also active in the case of PCD induced by abiotic stress, we subjected *Arabidopsis* seedlings, with and without PA pre-treatment, to NaCl stress. NaCl has previously been shown to induce a high rate of PCD in *Arabidopsis* root hairs (Kacprzyk et al., 2014), and pre-treatment with SPM and SPD at both 100 nM and 1 µM concentrations significantly inhibited NaCl-induced PCD (Figure 3) with root hairs remaining viable (Figure S2).

**Figure 3.**
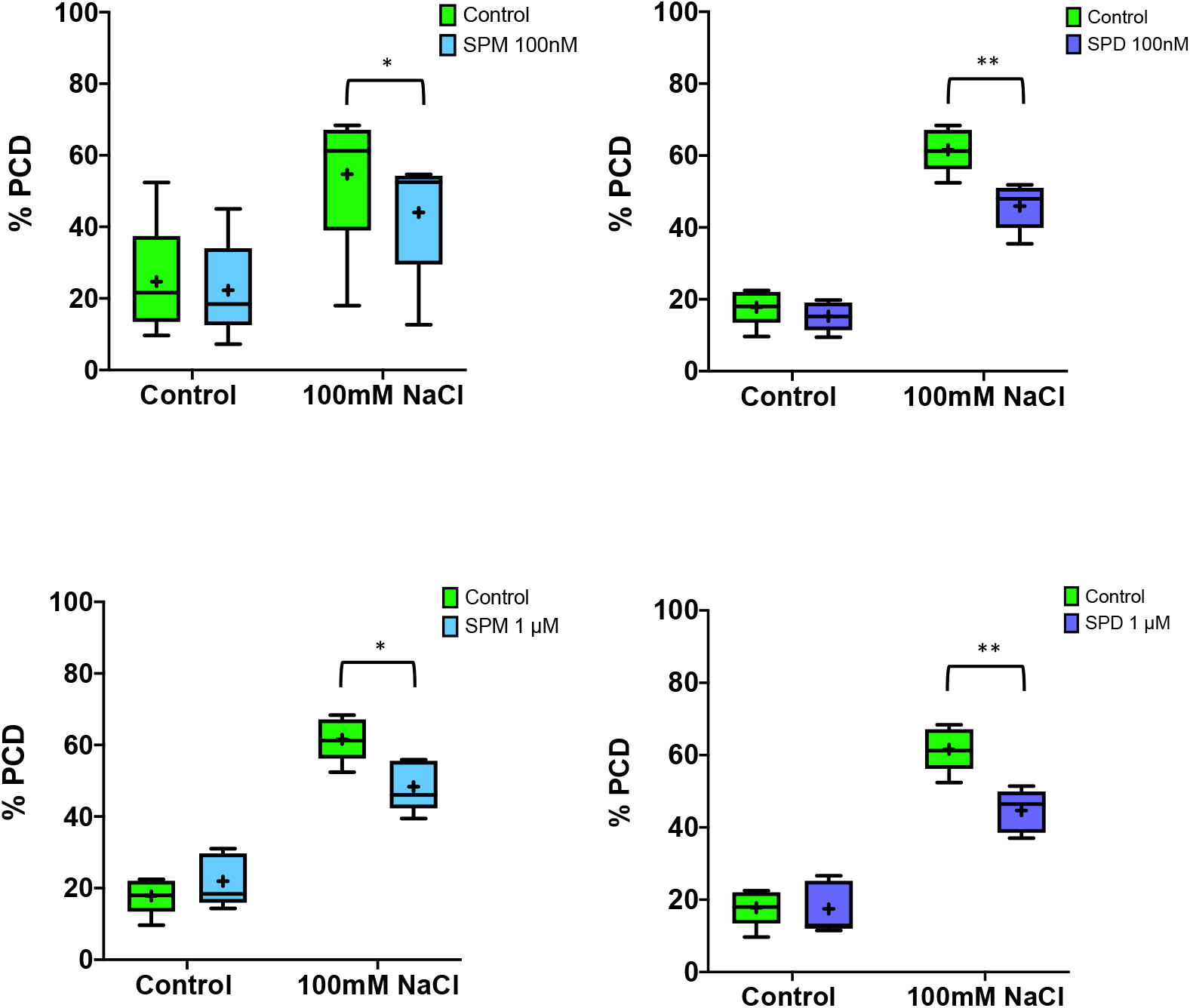
Pre-treatment with SPM or SPD protects *Arabidopsis* root hairs from NaCl-induced PCD. *Arabidopsis* seedlings were treated with 100 nM SPM, 100 nM SPD, 1 µM SPM or 1 µM SPD for 24 hours before treatment with 100 mM NaCl for 5 minutes. Root hairs were then scored as viable, PCD or necrotic 24 hours later. For each treatment, 2 seedlings were examined per experiment, and experiments were repeated 5 times. Rates of PCD root hairs were compared to control using 1-way ANOVA. *≤0.05, **≤0.01, ***≤0.001, ****≤0.0001. Lines on box and whisker plots show median values, crosses show mean values and whiskers show the minimum to maximum values.

### Treatment with low, PCD-inhibiting concentrations of spermine or spermidine causes a transient ROS burst in Arabidopsis roots, and induces expression of oxidative stress and pathogen responsive genes in ACSC*s*

Previous studies have demonstrated that catabolism of PAs can generate H_2_O_2_ in the context of PCD induction (Yoda et al., 2003; Yoda et al., 2006; Moschou et al., 2008b). To establish if the PA concentrations capable of inhibiting PCD in this study also induce increases in ROS levels, 5-day old *Arabidopsis* seedlings were treated with H_2_O (mock control), 1 µM SPM, 1 µM SPD or 1 mM H_2_O_2_ (positive control), and ROS levels in root tips were visualised by staining with H_2_DCFDA after 30 minutes (Figure 4A) and 2 hours (Figure 4B). Both PAs as well as H_2_O_2_ induced a significant increase in ROS in *Arabidopsis* root tips after 30 minutes, relative to control roots (Figure 4A). In SPM treated roots this increase in ROS levels was observed even after 2 hours of treatment, whereas ROS signal returned to control levels in H_2_O_2_ and SPD treated roots (Figure 4B). Furthermore, in order to establish if PCD-blocking concentrations of PAs are affecting transcriptional changes in cells, we carried out semi-quantitative PCR (semi-qPCR) on ACSC treated with mock H_2_O control, 1 µM SPM or 1 µM SPD at 3 time points; 2 hours, 6 hours and 24 hours post treatment. The expression of 4 marker genes for different stress responses was examined; *alternative oxidase 1a* (*AOX1a*, mitochondrial stress response marker and regulator of cell redox state (Van Aken et al., 2009)), *enhanced disease susceptibility 1* (*EDS1*, regulates hypersensitive response upstream of SA accumulation and downstream of ROS signalling (Rusterucci et al., 2001; Straus et al., 2010)), *glutathione S-transferase* (*GST1*, ROS-responsive antioxidant protein (Love et al., 2005)) and *pathogenesis related 1* (*PR1*, marker gene for SA dependent systemic acquired resistance (Gaffney et al., 1993; Cameron et al., 1999)). At 2 h post treatment, both exogenously applied PAs lead to increased expression of *AOX1a*, with the difference being statistically significant for SPM at 0.05 level (Figure 4C). Similarly, *PR1* expression was significantly increased by treatment with both PAs after 2 hours. Additionally, after 6 hours, SPM treatment also induced increased expression of *EDS1* (Figure 4C).

**Figure 4.**
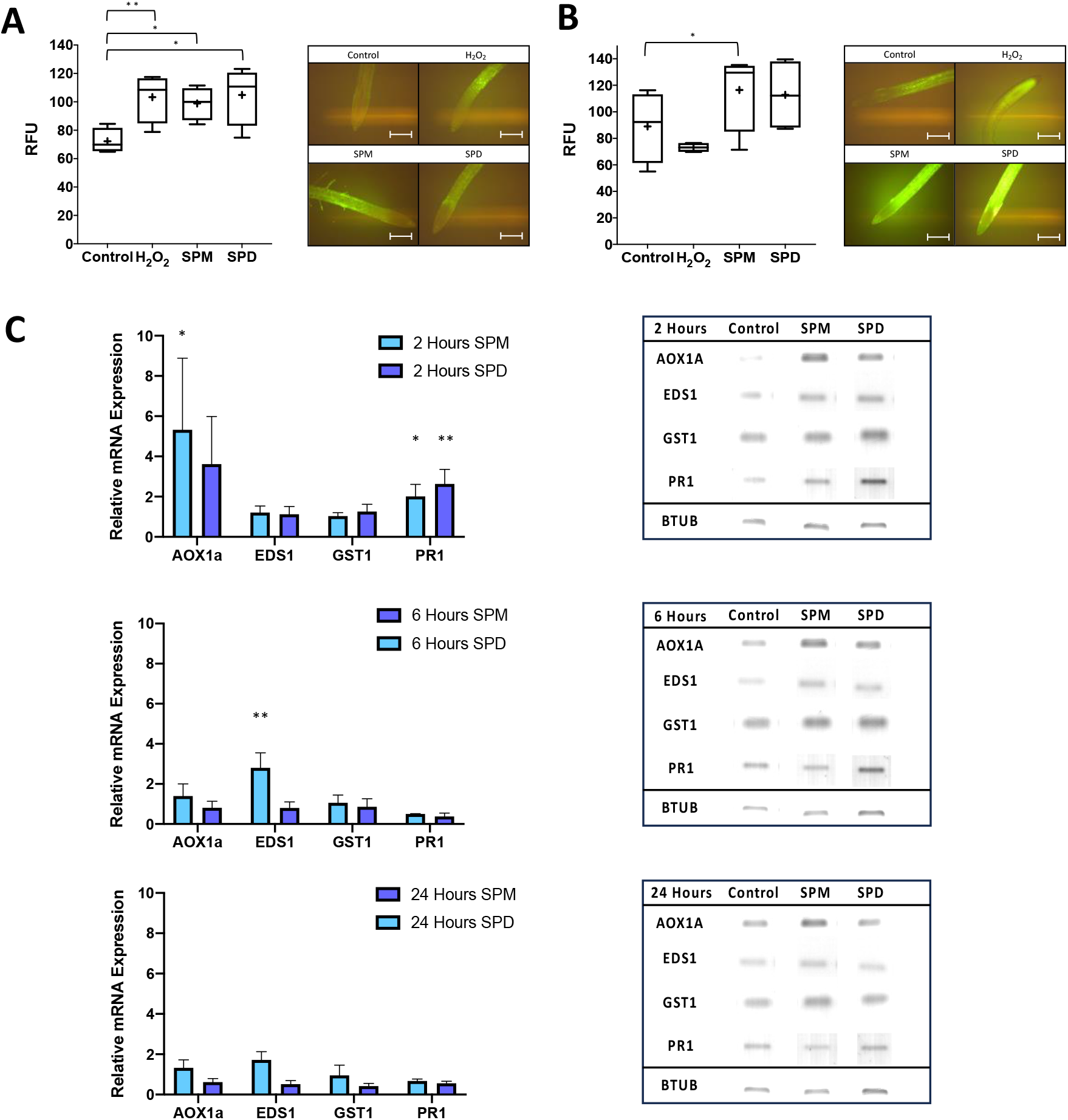
Low concentrations of SPM and SPD induce ROS bursts and lead to induction of ROS- and pathogen-responsive genes. 5-day old *Arabidopsis* seedlings were treated with H_2_O (mock control), 1 µM SPM, 1 µM SPD or 1 mM H_2_O_2_ for **(A)** 15 minutes or **(B)** 105 minutes and transferred to slides and incubated with H2DCFDA (10 µM) in the dark for a further 15 minutes before imaging. Seedlings imaged in the presence of H_2_O rather than H2DCFDA were used to establish baseline autofluorescence values, which were subtracted from experimental values for each replicate. 4-5 seedlings were imaged for each experiment, and experiments were repeated 4 times. Mean relative fluorescence units (RFU, measured as mean green pixel intensity) as well as representative images are shown for each timepoint. Lines on box and whisker plots show median values, crosses show mean values and whiskers show the minimum to maximum values. Scale bar = 200 µm. **(C)** 6-day old ACSCs were treated with mock H_2_O control, 1 µM SPM or 1 µM SPD. Cells were sampled for RNA-extraction 2 hours, 6 hours and 24 hours later and expression of *AOX1a*, *EDS1*, *GST1*, *PR1* determined using semi-quantitative PCD with *ß-tubulin* as the housekeeping control. ImageJ was used for densitometry, with band intensity of each gene normalised to ß-tubulin for each timepoint, and gene expression of SPM and SPD-treated samples presented as relative fold change over mock control. Experiments were carried out in triplicate, mean relative changes in gene expression for each timepoint are shown on the left, representative DNA gel images from the same experimental replicate are shown on the right. RFU for each treatment was compared to control using 1-way ANOVA. mRNA expression in treated samples were compared to control samples using 1-way ANOVA *≤0.05, **≤0.01, ***≤0.001, ****≤0.0001.

#### Stress sensitivity and PCD rates of *Arabidopsis* polyamine metabolism mutants

Null mutant plants deficient in synthesis of SPM (*spms*), thermo-spermine (*acl5*) and the *spms-acl5* double mutant have previously been used to characterise the roles of PAs in plant development and stress tolerance. Firstly, as previous studies reported increased sensitivity of *acl5* and *spms-acl5*, but not *spms*, to salinity at whole organism level (Yamaguchi et al., 2006; Alet et al., 2012) we tested if these findings also applied in our lines, measured as seedling survival (Figure 5A). The *acl5* and *spms-acl5* lines were more sensitive to oxidative stress as demonstrated by lower survival on medium supplemented with MV (Figure 5A), a ROS producing herbicide which has been shown to effect PA metabolism in plants (Kurepa et al., 1998; Fujita and Shinozaki, 2014). Subsequently, to explore whether these genotypes also show different responses to PCD inducing stimuli, we subjected 5-day old *spms*, *acl5* and *spms-acl5* seedlings to SA and NaCl treatments (Figure 5A). Root hairs were scored as viable, PCD or necrotic 24 hours post treatment. The *acl5* seedlings displayed lower PCD rates (Figure 5B) and higher viability (Figure S3) after SA treatment, but there were no other significant differences observed in root hair cell death.

**Figure 5.**
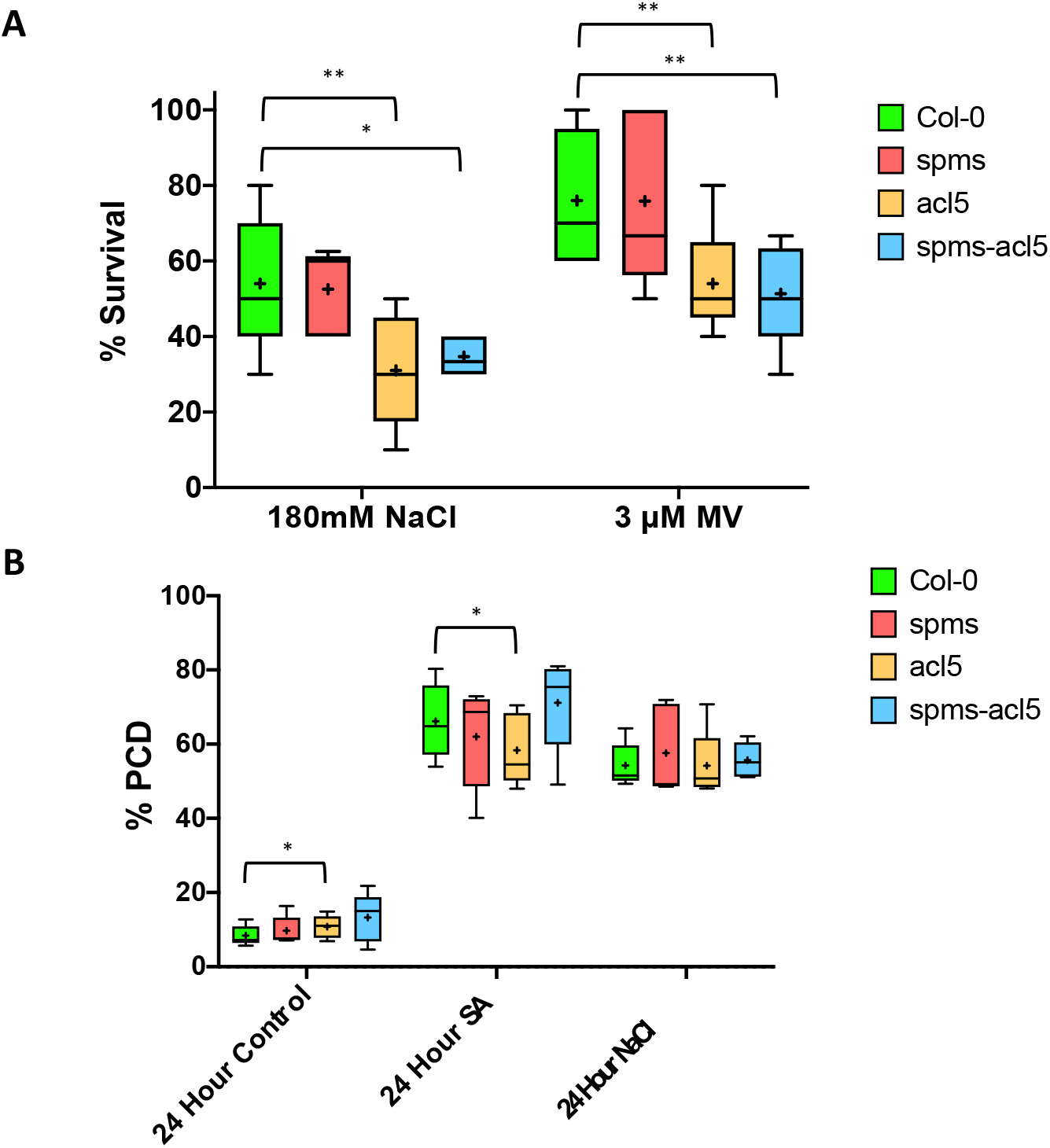
Seedling stress survival and root hair PCD rates of *Arabidopsis acl5*, *spms* and *spms-acl5* mutants. 7-day old *Arabidopsis* seedlings were transferred to agar plates containing 180 mM NaCl or 3 µM MV, and seedling survival scored after a further 7 days **(A).** For each treatment 10 seedlings per genotype were examined per experiment, and experiments were repeated 5 times. Survival rates in each treatment were compared to Col-0 using 1-way ANOVA. **(B)** 5-day old *Arabidopsis* seedlings were subjected to PCD induction with 20 µM SA and 100mM NaCl. Root hairs were then scored as viable, PCD or necrotic after 24 h, 2 seedlings were examined per treatment per experiment, and experiments were repeated 5 times. Rates of PCD in each treatment were compared to Col-0 control using paired 1-way ANOVA. *≤0.05, **≤0.01, ***≤0.001, ****≤0.0001. Lines on box and whisker plots show median PCD rates, crosses show means and whiskers show the minimum to maximum values.

## Discussion

The data presented herein provide new insights into the effect of treatment with exogenous PAs, and genetic modulation of PA metabolism, on the plant PCD response. We used two established models for quantitative determination of PCD rates *in vitro* (ACSC, Reape et al., 2008) and *in vivo* (root hair assay, Kacprzyk et al. 2015). ACSCs are widely used for studying plant PCD (McCabe and Leaver, 2000; Reape et al., 2008; Burke et al., 2023), as they facilitate cell death induction using a wide diversity of treatments or pharmacological manipulation of this process, and enables morphology-based monitoring of rates of cell death. The root hair assay (RHA) allows for quantitative determination of PCD response to stress treatments applied by examination of viability and morphology of root hair cells (Hogg et al., 2011), and provides a useful tool for testing the role of specific genes in PCD regulation as it allows determination of PCD rates in multiple mutant/transgenic lines in a relatively short time (Kacprzyk et al., 2014; Chua et al., 2020). We demonstrate that in both systems, treatment with two of the most abundant PAs in plants, SPM and SPD (Minocha et al., 2014; Jancewicz et al., 2016), induces viability loss, and development of hallmark PCD morphology (cytoplasmic condensation and protoplast shrinkage away from the cell walls (Kacprzyk et al., 2017)), that occurs over a time span similar to that of PCD induced by other chemicals and abiotic stimuli (<24 hours) (Reape et al., 2008). While both SPM and SPD have previously been shown to cause tissue damage in tobacco leaves, this damage was quantified only as “cell collapse” (Yoda et al., 2003) without examination of morphological features of the dying cells. PCD induction by exogenous PAs is in line with the observations from numerous experimental systems that report accumulation of PAs prior to PCD induction in plant cells and tissues. For example, PA accumulation was observed prior to PCD induction by cadmium treatment in both *Malus hupehensis* seedlings (Jiang et al., 2012) and tobacco BY-2 cells (Kuthanová et al., 2004), as well as during HR PCD in *Nicotiana tabacum* (Torrigiani et al., 1997; Yoda et al., 2006), rice and *Arabidopsis* (Yoda et al., 2009). However, the literature reports on effect of PAs on PCD are often conflicting, with many studies reporting pro-survival effect of exogenous PAs supplementation under conditions such as aluminium stress (Yu et al., 2018) and salinity stress (Duan et al., 2008), and increases in PAs levels are generally associated with both abiotic and biotic stress resistance (Hussain et al., 2011; Shao et al., 2022). To ascertain if exogenous PAs can indeed modulate PCD response in a concentration dependent manner, we tested the effect of pre-treatment with sublethal levels of SPD and SPM on cell death induced by SA, a phytohormone implicated in PCD induced by biotic stress (Alvarez, 2000; Raffaele et al., 2006; Radojičić et al., 2018). Both PAs inhibited SA-mediated PCD induction and viability loss in ACSCs and *Arabidopsis* root hairs. A similar protective effect was observed for pre-treatment with SPM and SPD in context of PCD induced by NaCl, suggesting it is not specific to cell death inducing stimuli and may operate in the context of PCD induced by both abiotic and biotic triggers.

PAs were recently suggested to elicit defence responses dependent on ROS signalling, including transcriptional reprogramming, and local accumulation of SA (Liu et al., 2020), and Pas degradation yielding hydrogen peroxide was suggested to contribute to the oxidative burst during HR associated PCD (Yoda et al., 2003). Likewise, PAs have long been associated with salt tolerance in crop species (Krishnamurthy and Bhagwat, 1989), with the proposed molecular mechanisms likely depending on their ability to modulate both redox homeostasis and ion transport (Saha et al., 2015). Furthermore, salt induced PCD in plant cells is dependent on ROS generation by NADPH oxidase (Monetti et al., 2014), in tobacco cells salt treatment was shown to trigger the release of SPM into the apoplast where its catabolism releases H_2_O_2_ upstream of PCD (Moschou et al., 2008b). Therefore, there is strong evidence for PAs playing a role in regulation of redox signalling in both abiotic and biotic stress contexts. As moderate levels of H_2_O_2_ induce transcription of pro-survival antioxidant genes but higher H_2_O_2_ concentrations can promote induction of PCD (Gechev et al., 2002), it is plausible that the dose dependent effect of exogenous PAs on cell death and survival is at least partially mediated through transient modulation of ROS signalling. ROS are key regulators of PCD in plants, and changes in ROS levels can shift cell fate towards death or survival depending on the strength and timing of the ROS signal (Van Breusegem and Dat, 2006; De Pinto et al., 2006). Conversion of SPM to SPD, and SPD to PUT is catalysed by PAO family enzymes, of which five have been identified in *Arabidopsis* (Moschou et al., 2008c). Both of these reactions yield H_2_O_2_, making PA catabolism a significant source of cellular ROS (Murray Stewart et al., 2018). To probe the effect of exogenous PAs application on the ROS signalling, we first used the fluorescent stain H_2_DCFDA that is an established tool for visualising ROS in plant roots (Shin et al., 2005; Ha et al., 2018). Low concentrations of SPD and SPM, that was shown to inhibit PCD induced by SA and NaCl, triggered ROS production in root tips after 30 minutes comparable to that induced by 1 mM H_2_O_2_. After 2 hours, H_2_O_2_ treated roots no longer displayed elevated ROS compared to control levels indicating successful detoxification by plant tissues. Interestingly, a statistically significant increase in ROS level was observed at this time point for SPM but not SPD, supporting the idea that the ROS signals observed were due to the sequential catabolism of SPM to SPD to PUT by PAO. The effect of exogenous PAs on ROS signalling is also supported by gene expression changes induced by SPM and SPD treatment in ACSCs. We used semi-quantitative PCR to measure the effect of exogenous PAs on the expression of marker genes for oxidative stress response (*AOX1A*, *GST1*) and pathogen response (*EDS1*, *PR1*) after 2, 6 and 24 h of treatment. The results suggested induction of *AOX1a* and *PR1* at the earliest time point tested (2h) by both PAs, and induction of *EDS1* after 6h by SPM. *AOX1a* encodes *alternative oxidase 1a*, a stress responsive mitochondrial protein that enables uncoupling of respiration and electron transport from ATP generation, allowing the cell to balance carbon and energy metabolism as well as redox state (Vanlerberghe, 2013). *AOX1a* also acts as a modulator of PCD, with changes in *AOX1a* expression altering the stress threshold at which PCD is triggered (Van Aken et al., 2009). Previously, it was demonstrated that exogenous PAs trigger an increase in oxygen consumption in *Arabidopsis* that is dependent on both ROS signalling and the AOX pathway (Andronis et al., 2014). The hypothesis that activation of *AOX1a* expression is responsible for the PCD-inhibiting effects of these PAs is supported by our finding that the expression of GST1, a general detoxifying enzyme that is responsive to various stressors (Wagner et al., 2002) was unchanged by treatment with exogenous PAs. This would suggest that PAs are not exerting this effect solely through the upregulation of plant antioxidant enzymes in response to increased cellular H_2_O_2_ levels. Further, *PR1* is the classic marker gene for SA accumulation and biotic stress in plants (van Loon et al., 2006) and exogenous PAs were previously shown to induce *PR1* expression in tobacco (Yamakawa et al., 1998). The increase in *PR1* expression observed in ACSCs suggested that PAs treatment lead to SA accumulation and activation of biotic stress responses in our model system, similarly to observations in *Arabidopsis* seedlings (Liu et al., 2020) that could explain their protective effect against SA induced PCD. The observed upregulation of EDS1 after 6h of SPM treatment also supports this observation. *EDS1* is a signalling protein involved in various aspects of plant defence, and is responsible for SA accumulation and the downstream transcription of canonical defence genes, including *PR1* (Feys et al., 2001), as well as the regulation of cell death during effector triggered immunity (Dongus and Parker, 2021). *EDS1*, with its co-receptor *PAD4*, also regulates the expression of *AOX1a* in the context of PCD induced by pathogen attack (Ho et al., 2008). Collectively, the obtained results suggest that low levels of exogenous PAs may inhibit PCD by triggering a transient ROS burst that primes the cell against future biotic or abiotic stress via transcriptional reprogramming.

To further explore the role of PA metabolism in PCD regulation, we subsequently investigated the cell death response in *Arabidopsis* plants deficient in the synthesis of SPM (*spms*), thermospermine (*acl5*) and the double mutant (*spms-acl5*) (Imai et al., 2004). The *spms* plants have reduced SPM levels but no obvious morphological phenotype under normal growth conditions (Imai et al., 2004), while *acl5* and *spms-acl5* display reduced stem growth (Kakehi et al., 2008) and xylogenesis defects (Muñiz et al., 2008). The whole seedling survival experiments we performed indicated that both *acl5* and *spsms-acl5* seedlings are more sensitive to long term salinity and herbicide (methyl viologen) induced oxidative stress. The salinity tolerance phenotype was previously reported for these lines (Yamaguchi et al., 2006; Alet et al., 2012). While exogenous PAs supplementation increased MV tolerance in *Arabidopsis* (Kurepa et al., 1998), sunflower (Benavides et al., 2000) and tomato (Pascual et al., 2023), increased sensitivity of thermospermine synthesis mutants to this widely used herbicide has not been previously reported. Genetic engineering focused on increasing the endogenous levels of this PA thus suggesting a possible route by which herbicide resistance in crop species could be improved, provided that negative developmental consequences can be avoided. However, when we tested the rates of SA and NaCl-induced cell death in the root hairs of *spsms*, *acl5* and *spsms-acl5* seedlings, the only detected phenotype was a minor decrease (8%) in rates of SA-induced PCD observed for the *acl5* mutant. Of course, it needs to be noted that while root hair assay allows rapid screening of mutant genotypes (Hogg et al., 2011), it is a method based on observation of stress induced PCD in just one cell type. This observation also emphasises that there are numerous, likely antagonistic, mechanisms via which PA metabolism can modulate plant stress responses and PCD which might be difficult to dissect in mutant backgrounds, especially in the case of lines showing developmental defects such as *acl5*. Indeed, despite exogenous supplementation with SPM appearing to induce transient increase in ROS production here and in other studies (Yoda et al., 2003; Moschou et al., 2008b), the *spms* mutant that is deficient in SPM synthesis has been recently shown to have an enhanced flg22-triggered ROS burst (Zhang et al., 2023). Additionally, while the same study also observed an increase in ROS following treatment with low concentrations of SPM alone, co-treatment or pre-treatment with SPM inhibited the flg22-triggered ROS burst. Further pharmacological and genetic dissection of the role of SPM in this context led the authors to suggest that SPM was exerting this effect on ROS production through direct inhibition of *respiratory burst oxidase homologue D* (RBOHD) inhibition (Zhang et al., 2023). This evidence, combined with the fact that spermine has been shown to act directly as a scavenger of ROS (Ha et al., 1998), highlights the complex and numerous molecular functions of PAs that makes it difficult to categorize them as either pro-survival or cell death promoting. In conclusion, here we demonstrate that at higher concentrations exogenous SPM and SPD can induce PCD characterised by the hallmark morphology of retracted protoplast in *Arabidopsis,* whereas pretreatment with lower concentration of these PAs can inhibit PCD included by SA and NaCl, at least partially through ROS mediated transcriptional reprogramming. However, further studies are required to fully elucidate the complex relationship between PA metabolism and PCD regulation.

## Acknowledgements

We wish to thank University College Dublin and the School of Biology an Environmental Science for funding the PhD project of Rory Burke under the PhD Advance Scheme. Daniele Nicotra’s work was funded by an Erasmus Traineeship.

## Materials and Methods

### Plant Material

ACSC (ecotype *Ler*) was maintained as described in (May and Leaver, 1993), and sub-cultured weekly by transferring 10 mL of 7-day cell culture to 100 mL of fresh media. Cells grown in the dark for 6 days at a 21°C were used for experiments. *Arabidopsis thaliana* mutants (Col-0 background) in genes mediating PA metabolism (*spms-1*, *acl5-1, acl5-1 spms-1*) (Imai et al., 2004) were kindly provided by Professor Taku Takahashi. Seeds were sterilized and plants grown as previously described (Hogg et al., 2011) on half-strength Murashige and Skoog medium (2.15 g/L MS basal salts, 1% sucrose, pH 5.8) solidified with 1% agar in square Petri dishes at 21°C (16 h light/8 h dark).

### PCD induction in ACSC

For PA toxicity experiments, 20 mL aliquots of 6-day old cells were aseptically transferred to 100 mL Erlenmeyer flasks and treated with indicated concentrations of SPM and SPD (Sigma). Stock solutions (1M) of SPM and SPD were prepared in deionised water. For salicylic acid (SA) experiments, 6-day old cells were pre-treated with PAs or H_2_O control, before addition of SA (1.5 mM), (Merck, Burlington, MA, USA) or solvent control (0.1% ethanol) 24 hours later.

### PCD induction in root hairs

For PA toxicity experiments, 5-day old seedlings were transferred to 1 mL of sterile water in a 24 well plate containing 10 µM, 100 µM or 1mM SPM or SPD. Seedlings were returned to standard growth conditions for 24 h before scoring. For SA experiments, 4-day old seedlings were pre-treated with 100 nM or 1 µM SPM/SPD for 24 h, then treated with 20 µM SA and returned again to standard growth conditions until scoring for cell death rates. For NaCl experiments, 4-day old Col-0 seedlings were pre-treated with PAs, submerged in 100mM NaCl solution for 5 minutes (or water for control treatment) and returned to 1 mL SDW until scoring for PCD rates.

### Determination of PCD, viability and necrosis rates in ACSC and root hairs

ASCSs and root hairs, incubated in the viability stain fluorescein diacetate (FDA), were scored using light microscopy as previously described (Kacprzyk et al., 2017). Briefly, cells/root hairs positive showing green fluorescence after FDA staining were scored as viable, while dead cells/root hairs were scored as either PCD or necrotic based on the presence or absence of protoplast retraction (hallmark PCD morphology) respectively. For suspension cells, at least 200 cells were scored per treatment. For root hairs, 2-3 seedlings (approx. 150-200 root hairs each) were scored per condition/treatment.

### Seedling Survival Assay

The seedling survival assay was performed as previously described (Burke et al., 2023). Briefly, 7-day old seedlings were transferred aseptically to square petri dishes containing half-strength Murashige and Skoog medium (2.15 g/L MS basal salts, 1% sucrose, pH 5.8) solidified with 0.6% agar and supplemented with either 180 mM NaCl or 3 µM methyl viologen (MV) (ThermoScientific). Seedlings were then grown horizontally under standard growth conditions for a further 7 days before being categorized as viable or dead, based on bleaching of the newest emerging leaves.

### ROS specific staining and quantification of Arabidopsis Root Tips

Five-day old Col-0 seedlings were transferred to wells of a 24-multiwell plate containing 1 mL of sterile water (control), SPM (1 µM), SPD (1 µM) or H_2_O_2_ (1 mM) (Sigma). For ROS visualisation seedlings were stained, at indicated time points, directly on a microscope slide with 10 µM H_2_DCFDA (Thermo) or sterile H_2_O (blank control) and incubated in the dark for a further 15 minutes before imaging using a Leica DM500 fluorescence microscope (excitation 475 nm, emission 510 nm). For each treatment 5 seedlings were imaged, and the experiment was repeated 4 times. Captured images were imported to ImageJ (Schneider et al., 2012), root tip area was selected manually, and mean green pixel intensity for the root tip area calculated using the ‘RGB Measure’ analysis plugin. The mean ‘blank’ pixel intensity (H_2_O control, no H2DCFDA) for each experiment was then used to normalise the values for each experimental replicate.

### RNA Isolation

Twenty mL aliquots of 6-day old *Arabidopsis* dark grown cell cultures were transferred to 100 mL Erlenmeyer flasks and treated with SPM (1 mM), SPD (1 mM) or solvent control (H_2_O) in the dark. At 2 hours, 6 hours and 24 hours post treatment, 5 mL of cells were withdrawn from each flask, medium removed using vacuum filtration through Whatman grade 1 paper and cells flash frozen using liquid nitrogen in 2 mL tubes containing ∼20 chrome/steel beads and stored at −80°C prior to RNA extraction. Frozen cells were homogenized using a mixer mill (Retsch MM400). RNA isolation was carried out using a QIAGEN RNeasy Plant Mini kit, including the on-column DNAse treatment step. Total RNA obtained for each sample was quantified using Nanodrop (Thermo Scientific).

### Semi-quantitative PCR

cDNA synthesis was carried out using the ThermoScientific first-strand cDNA synthesis kit according to the manufacturer’s instruction with 1 µg of total RNA used per each reaction. Semi-quantitative PCR was carried out using the DreamTaq PCR mastermix (ThermoScientific). Primers and reaction conditions used are described in Table S1. PCR products were run on ethidium bromide supplemented 1.5% agarose gel at 80 volts. Imaged gels were imported to ImageJ (Schneider et al., 2012) for densitometry analysis. Mean band intensity for each product was normalized to BTUB for each timepoint.

### Statistical Analysis

Statistical tests were carried out using GraphPad Prism version 8.2.1 for Mac OS (GraphPad Software, San Diego, CA, USA). For cell culture and root hair assay experiments, paired one-way ANOVA tests without multiple testing correction were used to compare viability and PCD rates between treatments. For survival and RHA experiments involving PA metabolism mutant lines, paired one-way ANOVA tests without multiple testing correction were used to compare survival or viability and PCD rates respectively for the same treatment across genotypes.

## Supplemental Tables

**Table S1.**
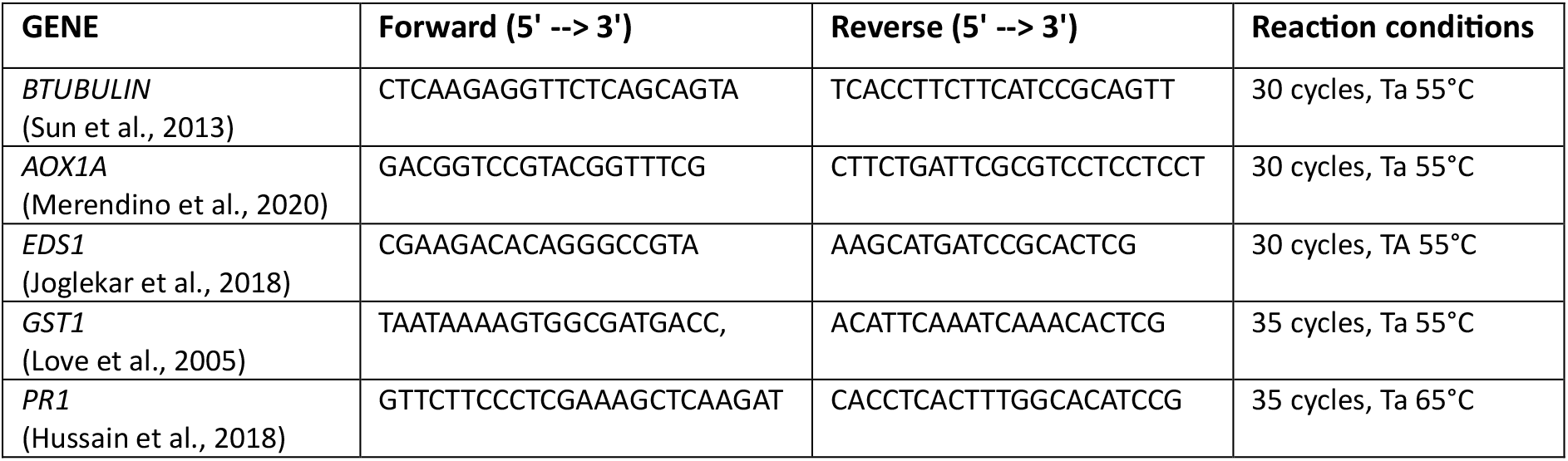
Primers used in this study.

## Supplemental Figures

**Figure S1.**
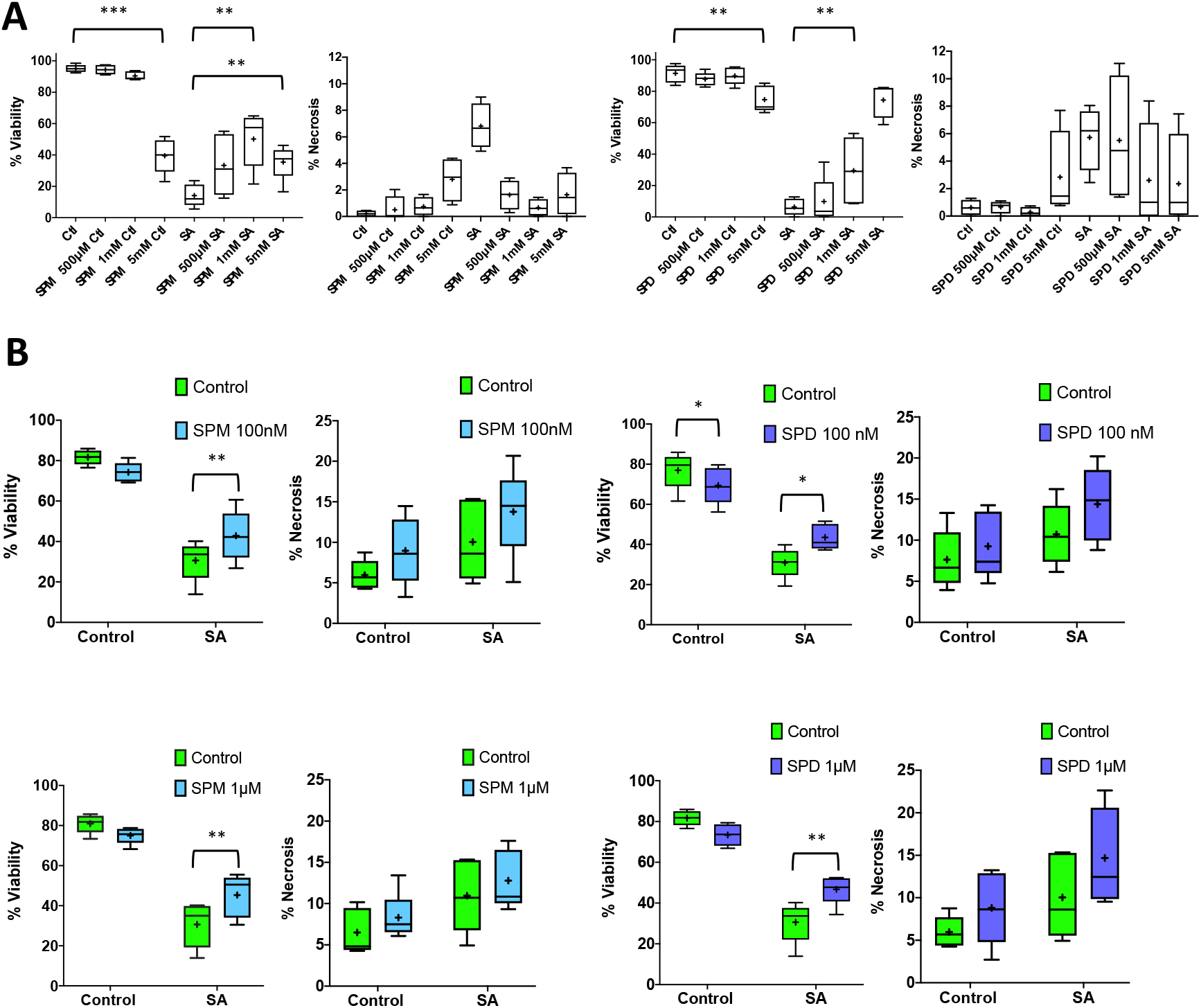
Pre-treatment with SPM or SPD maintains viability in ACSCs and *Arabidopsis* root hairs treated with SA. ACSCs were treated with indicated concentrations of SPM or SPD for 24 h prior to PCD induction with 1.5mM SA **(A)**. Cells were scored as viable, PCD or necrotic 24 h later. Experiments were repeated 5 times and rates of viable and necrotic cells in each treatment compared to control using 1-way ANOVA. *Arabidopsis* seedlings were treated with 100 nM SPM, 100 nM SPD, 1 µM SPM or 1 µM SPD for 24 hours, before PCD induction with 20 µM SA **(B)**. Root hairs were then scored as viable, PCD or necrotic 24 hours later. For each treatment 3 seedlings were examined per experiment, and experiments were repeated 5 times. Rates of viable and necrotic root hairs in each treatment were compared to control using 1-way ANOVA. *≤0.05, **≤0.01, ***≤0.001, ****≤0.0001. Lines on box and whisker plots show median values, crosses show mean values and whiskers show the minimum to maximum values.

**Figure S2.**
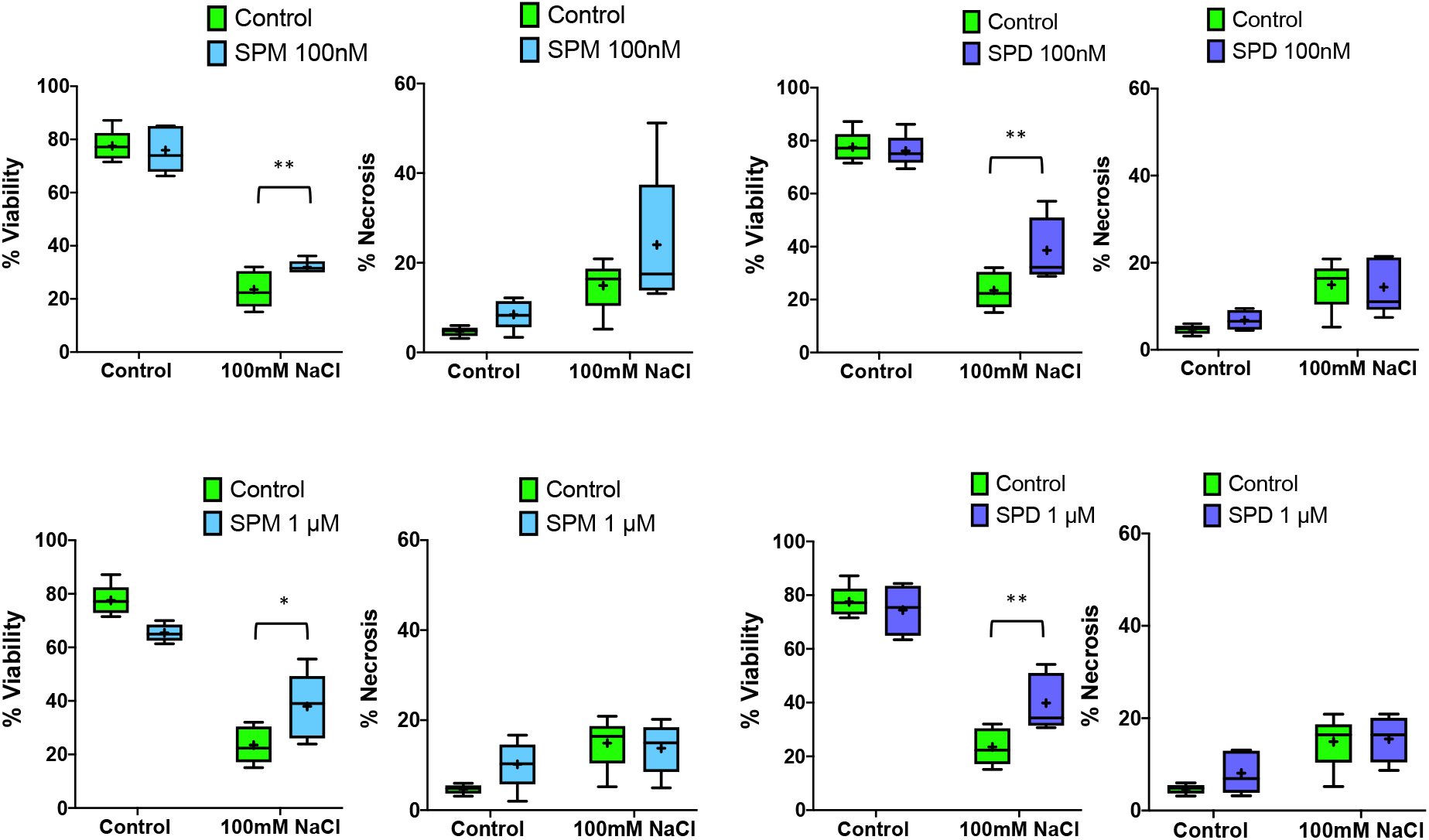
Pre-treatment with SPM or SPD maintains viability in *Arabidopsis* root hairs treated with NaCl. *Arabidopsis* seedlings were treated with 100 nM SPM, 100 nM SPD, 1 µM SPM or 1 µM SPD for 24 hours before treatment with 100 mM NaCl for 5 minutes. Root hairs were then scored as viable, PCD or necrotic 24 hours later. For each treatment, 2 seedlings were examined per experiment, and experiments were repeated 5 times. Rates of viable and necrotic root hairs were compared to control using 1-way ANOVA. *≤0.05, **≤0.01, ***≤0.001, ****≤0.0001. Lines on box and whisker plots show median values, crosses show mean values and whiskers show the minimum to maximum values.

**Figure S3.**
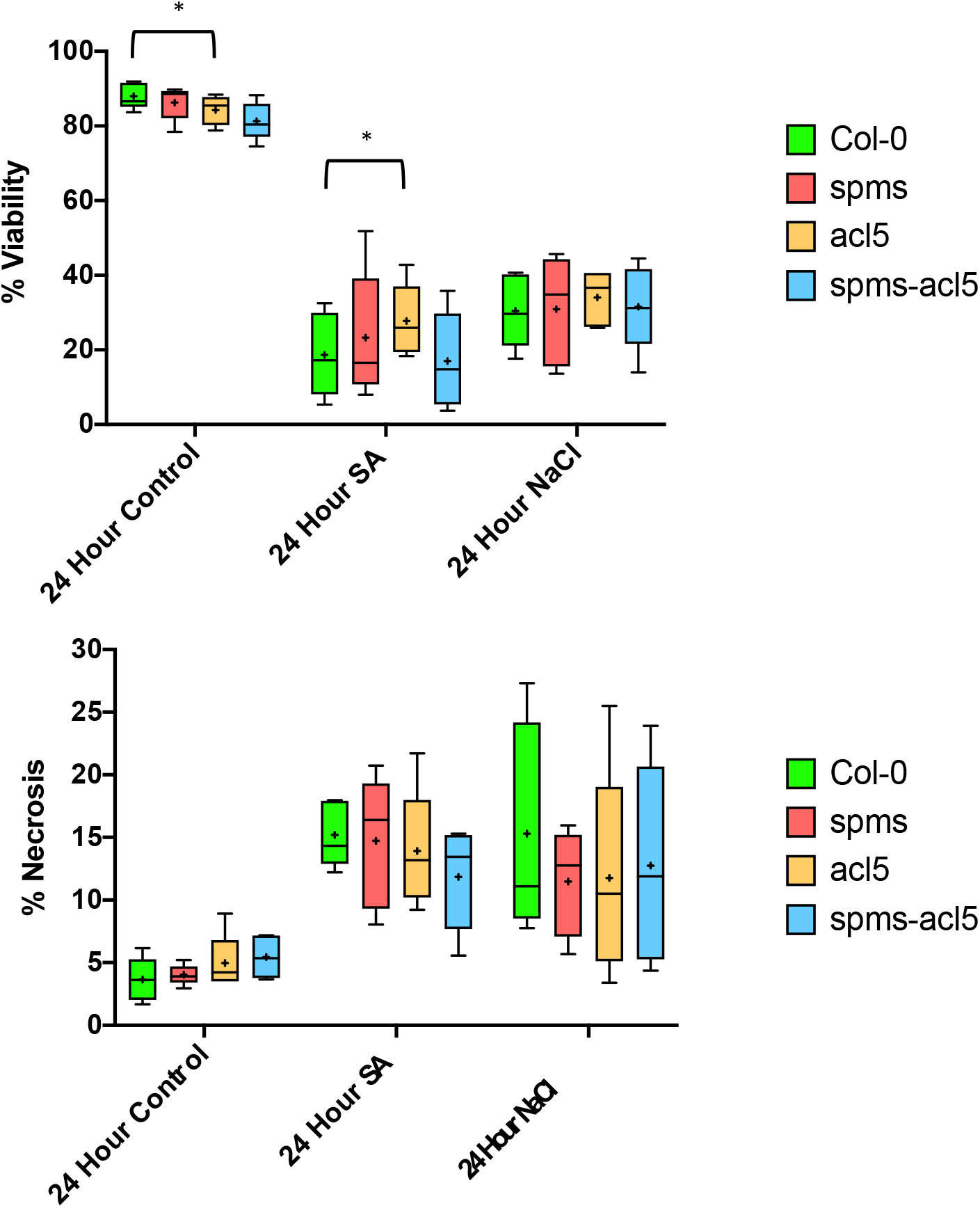
Root hair viability and necrosis rates of *Arabidopsis acl5*, *spms* and *spms-acl5* mutants. 5-day old *Arabidopsis* seedlings were subjected to PCD induction with 20 µM SA and 100mM NaCl. Root hairs were then scored as viable, PCD or necrotic after 24 h, 2 seedlings were examined per treatment per experiment, and experiments were repeated 5 times. Rates of viability and necrosis in each treatment were compared to Col-0 control using paired 1-way ANOVA without multiple testing correction. *≤0.05, **≤0.01, ***≤0.001, ****≤0.0001. Lines on box and whisker plots show median PCD rates, crosses show mean and whiskers show the minimum to maximum values.

